# A model-free method for genealogical inference without phasing and its application for topology weighting

**DOI:** 10.1101/2025.07.22.666161

**Authors:** Simon H. Martin

## Abstract

Recent advances in methods to infer and analyse ancestral recombination graphs (ARGs) are providing powerful new insights in evolutionary biology and beyond. Existing inference approaches tend to be designed for use with fully-phased datasets, and some rely on model assumptions about demography and recombination rate. Here I describe a simple model-free approach for genealogical inference along the genome from unphased genotype data called Sequential Tree Inference by Collecting Compatible Sites (sticcs). sticcs applies a heuristic algorithm based on the perfect phylogeny principle to reconstruct a local genealogy at each variant site in the genome, using a ‘collecting’ procedure to import information from other nearby sites. Using simulations, I show that sticcs is accurate for ARG inference, but only when the sample size is small. However, I also describe how it can be applied for the purpose of topology weighting by ‘stacking’ tree sequences inferred for multiple subsets of the data, removing the sample size restriction. Topology weights estimated in this way from unphased data tend to be more accurate than those computed with full ARGs inferred from perfectly phased data using several popular tools. The methods presented therefore have promise for analysis of relatedness and introgression in non-model species, including polyploids. The new methods are implemented in two Python packages, sticcs (for ARG inference) and twisst2 (for topology weighting using the stacking procedure), which both interface with the tskit library for analysis of tree sequence objects.

## INTRODUCTION

The genealogical history of a set of samples and how it varies along the genome due to recombination (i.e. the ancestral recombination graph or ARG) carries rich information about population history and evolutionary processes like selection and gene flow. Recently, there have been major advances in methods to infer the ARG from genome sequence data (Rasmussen et al. 2014; Kelleher et al. 2019; Speidel et al. 2021; B.C. Zhang et al. 2023; Deng et al. 2024), along with efficient data structures such as the succinct tree sequence (Kelleher et al. 2018; Kelleher et al. 2019; Wong et al. 2024) and methods to analyse and summarise these data structure (Ralph et al. 2020; Speidel et al. 2021; Whitehouse et al. 2024). Topology weighting is one such method that summarises complex genealogical relationships among a defined set of groups that are not necessarily reciprocally monophyletic (e.g. species, populations or ecotypes) (Martin and Van Belleghem 2017). Given a genealogy in which each of the defined groups is represented by one or more tips, topology weights represent the proportion of subtrees with a particular topology, considering only subtrees in which each group is represented by a single tip. Since the number of possible bifurcating tree topologies for a small number of groups is manageable (three ingroups have three rooted topologies, four ingroups have fifteen etc.), the relationships are summarised by a small number of values that sum to 1. Topology weighting has been used to study species barriers (Martin et al. 2019), to investigate targets of adaptive introgression (Marburger et al. 2019), and even to identify genomic regions associated with trait variation (Belleghem et al. 2017; Stankowski et al. 2023).

The publication that described topology weighting used a crude approach to infer the input genealogies: simply divide the genome into narrow non-overlapping windows and infer local trees for each window (e.g. using neighbour joining) (Martin and Van Belleghem 2017). Windows of 50 SNPs were proposed as a compromise between power and resolution. This approach is obviously sub-optimal, not only because it does not attempt to identify the true breakpoints between genealogies along the genome, but also because it fails to exploit the fact that nearby regions of the genome tend to have very similar genealogies, and therefore variants outside of a focal interval can be informative about the genealogy at the focal interval. Numerous more advanced approaches for ARG inference have been described see (Nielsen et al. 2025) for a comprehensive review. These address the shortcomings described above, and are therefore in principle able to infer resolved genealogies with greater power and resolution, even those spanning very narrow genomic ranges (at little as 1 bp). However, these methods have primarily focused on reconstructing genealogies for single species, and have not been tested extensively in the context of multi-species datasets in which underlying model assumptions about rates of coalescence and recombination may be violated.

One general feature of existing approaches for ARG inference is that the unit of interest is the sequence haplotype. The placement of each haplotype in each local genealogy depends on its similarity to other haplotypes in the dataset, or to inferred ancestral haplotypes. As such, most existing approaches require phased haplotypes as input. One exception is the fully probabilistic approach of Argweaver (Rasmussen et al. 2014), which can explore alternative phasing via Markov Chain Monte Carlo (MCMC) sampling, but is much slower than other modern methods (see Results for comparisons). For all other existing methods, unless direct haplotype phasing is possible (e.g. from long-read sequencing), probabilistic inference of phase from diploid genotypes is needed. This requires very large sample sizes to achieve acceptable accuracy, and also makes underlying model assumptions including panmixia, making phasing problematic for small, multi-species datasets.

Here I describe sticcs, a simple heuristic approach for ARG inference with minimal model assumptions and without the need for genotype phasing. Rather than inferring local genealogies and their breakpoints from sequence haplotypes, the proposed approach aims to infer the genealogy *at each variant site* using the principles of perfect phylogeny (Gusfield 1991), in which the unit of inference is the variant ‘pattern’ rather than the sequence haplotype. The general idea is that a tree is inferred at each variant site by ‘collecting’ additional information from surrounding sites. This is similar to the approach of RENT+ (Wu 2011; Mirzaei and Wu 2017), but sticcs extend the idea to infer trees from unphased genotypes of any ploidy level, including mixed ploidies. However, while the inferred trees include a tip (or leaf) for each haplotype in the dataset (i.e. two tips for each diploid, four from each tetraploid, etc.) tips from the same individual should be considered as interchangeable, and not as phased haplotypes. Such redundantly labelled tree sequences are nevertheless useful for various applications, including topology weighting. Using simulated data, I show that ARG inference using sticcs is comparable in accuracy to state-of-the-art methods, but only when sample size is very small. For the purpose of topology weighting, however, this limitation can be overcome by iteratively inferring ARGs for subsets of the full dataset and ‘stacking’ the results. This stacking approach proves to be accurate when applied to simulated data, and it is able to identify known sites of adaptive introgression in real data from *Heliconius* butterflies.

## METHODS

### Overview

ARG inference using sticcs involves two steps: (1) Collecting site patterns that are mutually ‘compatible’, and (2) building local genealogies from each collection using perfect phylogeny. Below, I describe these in reverse order: first the tree building approach using phased or unphased (diploid or higher ploidy) genotypes, and then the procedure used to ‘collect’ compatible sites for tree building. sticcs is available as Python package that can be used from the command line or via a Python API. The input is genotype data in the vcf format (phased or unphased), and output is a tree sequence for each chromosome in either tskit format or as a newick text file.

As shown in the Results, sticcs is accurate for small sample sizes. I therefore also describe a procedure for topology weighting by reconstructing multiple small sub-ARGs and ‘stacking’ the resulting topology counts, which is applicable to datasets of any size. The stacking approach is implemented by a second python package, twisst2, which can also be used for topology weighting from a pre-inferred ARG. twisst2 can also be used from the command line or via its Python API.

### Perfect phylogeny for any ploidy

Under the infinite-sites mutational model (no recurrent mutation), all sequences that share the derived allele at a given variant site must descend from a common ancestor. In this way, each unique variant site ‘pattern’ (i.e. the list indicating which sequences carry the derived allele at that site) represents a unique ‘node’ in the local genealogy (Fig. 1A). Given a set of site patterns that are ‘compatible’ (the nodes they represent can all be placed in a single tree), the tree can be directly inferred (Gusfield 1991) (e.g. Fig. 1A). This is the concept of perfect phylogeny.

**Fig. 1.**
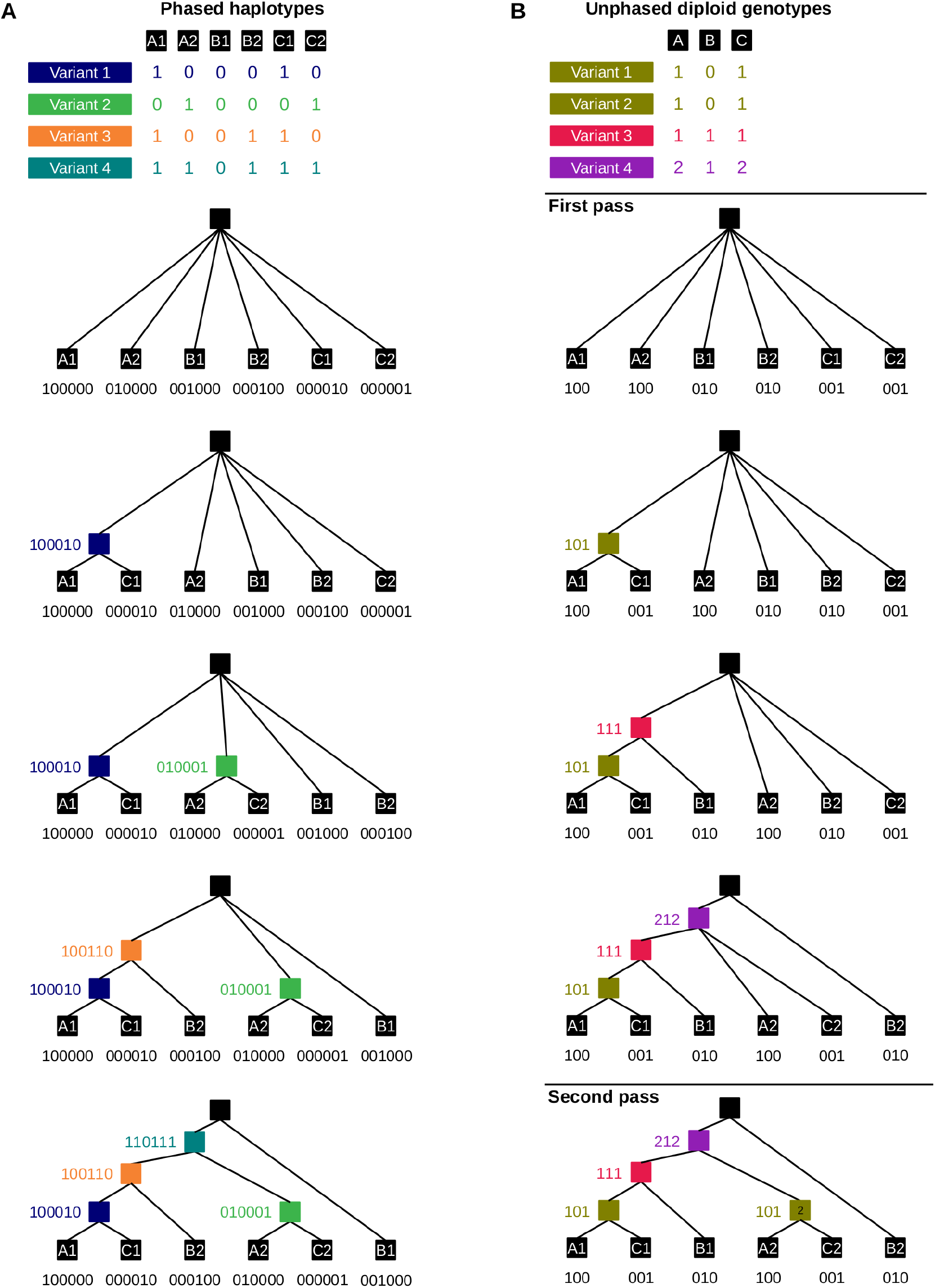
Tree building using perfect phylogeny for haploid and unphased diploid genotypes. Tables show variant patterns for four variants with three diploid individuals. Each pattern is the derived allele count for each haplotype (A) or unphased diploid genotype (B). The tree building procedure involves successively adding nodes to the tree such that the sum of the patterns of the child nodes is equal to the pattern of the parent node to be added. Note that with unphased diploid genotypes, the two leaf nodes from each individual have the same node pattern. This ambiguity results in a slightly different final tree (see leaves B1 and B2). In the diploid case, a second pass is included, in which a second node with pattern [1,0,1] is added.

The same logic can be extended to infer genealogies from unphased diploid (or higher ploidy) genotypes (Fig. 1B), as well as mixed ploidies. There are three important differences when applying perfect phylogeny to unphased data. First, site patterns no longer indicate which specific sequences carry the derived allele, but rather *how many* derived alleles are carried by each individual (0, 1 or 2, in the case of diploids). Patterns that include a 1 (indicating a heterozygote), correspond to nodes with one descendant (or tip) from the individual in question, but they do not tell us whether this tip should be the first or second haplotype. Hereafter, such patterns, and their corresponding nodes are referred to as ‘ambiguous’. Generalising to any ploidy, we can say that whenever the derived allele count is less than the full ploidy (but greater than zero) for any individual, the pattern is ambiguous. Phased or haploid data is then just a special case in which there are no ambiguous patterns. The most important consequence of the ambiguity of unphased data is that tips from the same individual (e.g. B1 and B2 in Fig. 1B), should be considered as interchangeable. For many downstream applications, such as topology weighting, this is inconsequential, because the two tips from each individual represent the same group/population.

The second difference when applying perfect phylogeny to diploids or higher ploidies is that the same site pattern may be shared by multiple nodes in a single tree. This occurs when all individuals carry either zero or one derived alleles at a site (i.e. nobody is homozygous derived). Consider for example variants 1 and 2 in Fig. 1. These two variants correspond to distinct nodes in the tree, but in diploid data they have the same pattern [1,0,1]. Hereafter, these are referred to as ‘duplicate nodes’. Patterns consistent with duplicate nodes are referred to as ‘duplicable’, because we cannot determine from the pattern alone how many nodes it corresponds to. Generalising to any ploidy, a pattern is duplicable when no individual carries more than half the full ploidy of derived alleles.

The final difference is that it is possible for a set of variant patterns to be mutually compatible with a single tree in all pairwise combinations, but for the complete set of variant patterns to be incompatible with a single tree. An example is shown in Fig. S1. This again is a consequence of the information lost in unphased data, which means that the test for pairwise compatibility (described in detail below) makes a maximum parsimony assumption. These three differences mean that the tree that is constructed can depend on the order in which nodes are incorporated. How the node order is decided in sticcs is described in the next section.

For now, assuming we have an ordered list of variant patterns, tree building in sticcs follows the following procedure (Fig. 1). The initial tree is unresolved, and comprises only a root node and 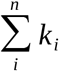 leaves, where *n* is the number of individuals and *k*_*i*_ is the ploidy of individual *i*.

Each leaf node is assigned a pattern consisting of n values, with 1 for the individual the leaf represents and 0 for all the other individuals. Only internal nodes (those that are not leaves or the root) can be added to the tree. These correspond to patterns where the total number of derived alleles is at least two and less than *n*. To add the first node, we attempt to pick its children from the available leaf nodes. Considering each leaf in order, we retain the leaf as a child node if each element in its pattern is ≤ the corresponding element in the parent node pattern. Subsequent children are added if the element-wise sum of child node patterns remains ≤ the parent node pattern. For example, node [1,0,1] has child nodes [1,0,0] and [0,0,1]. Any available child node that would cause the element-wise sum of child node patterns to exceed the parent node pattern is skipped. Once the element-wise sum of child node patterns is equal to the parent node, a suitable set of children has been found and the new internal node is incorporated into the tree. Subsequent internal nodes are added to the tree following the same procedure, except that previously-added internal nodes can also be children of the new node. With unphased data, the addition of a pattern may fail (a suitable set of children is not found, see Fig. S1), and the pattern is discarded. Once all site patterns have been added (or failed to be added), if the tree is not completely resolved (i.e. it contains at least one polytomy) the algorithm by default performs a second pass (or multiple passes for higher ploidies), in which it attempts to re-add nodes for any duplicable patterns (Fig. 2B).

**Fig. 2.**
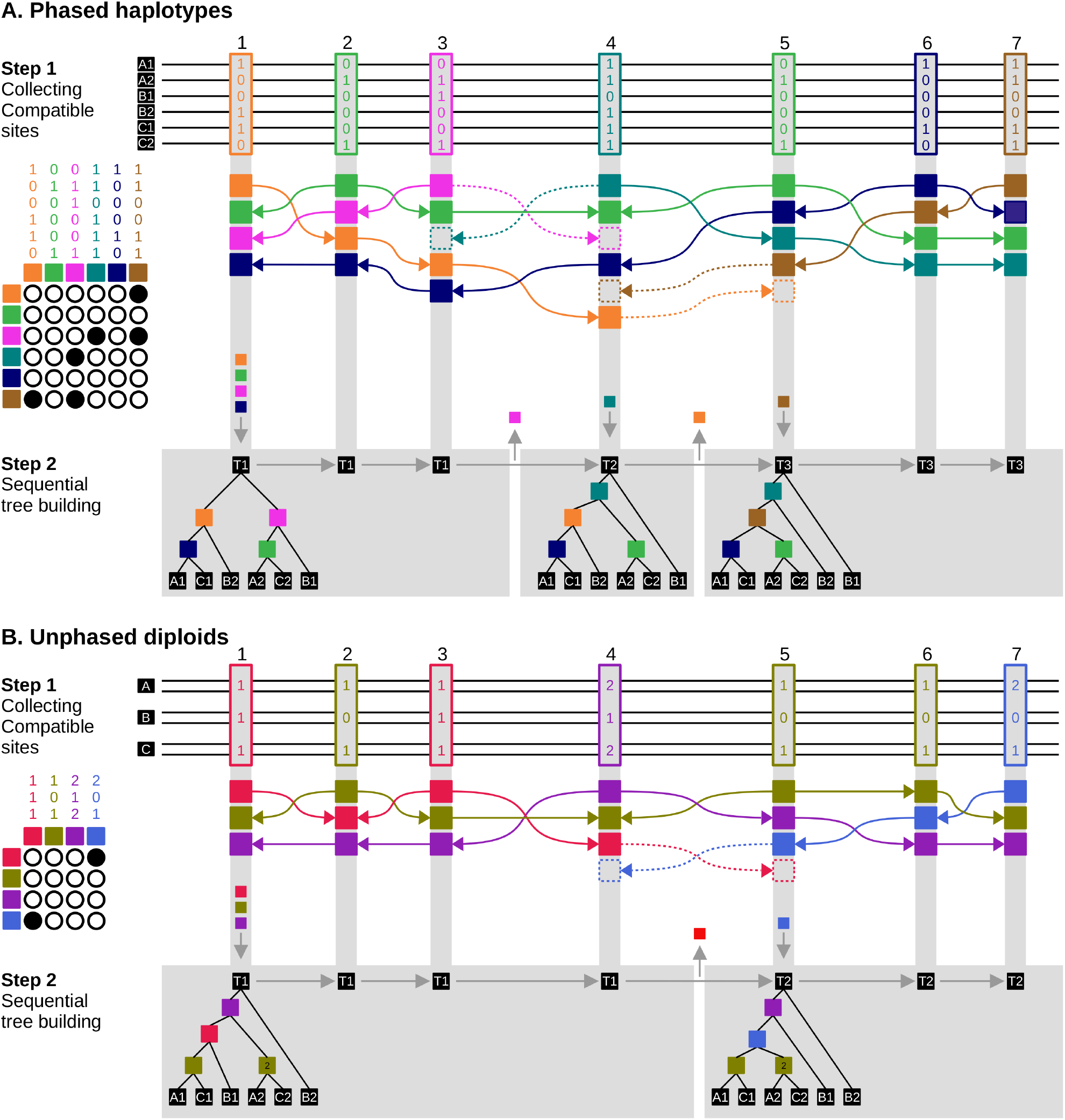
The sticcs algorithm applied to phased (A) and unphased (B) genotypes. In both panels genotypes from three diploid individuals are shown, either as six phased haplotypes (A) or three unphased genotypes (B). Each of the seven variant sites is given as a ‘pattern’ indicating the number of derived alleles (0 or 1 for phased data; 0, 1 or 2 for unphased). For each site, compatible patterns are collected from the left and right in order of distance (Step 1). Pairwise compatibility is indicated by open circles in the grid on the left. Patterns that are incompatible with the focal pattern at a given site are ‘knocked out’ (indicated by dashed lines and boxes) and become unavailable for collection beyond that site, but they may be considered for collection again under the ‘second chances’ rule described in the section on collecting compatible sites in the Methods. Tree building (Step 2) for each site follows the procedure shown in Fig. 1. Tree building for the second tree and onward is made more efficient by leaving in place nodes that are shared between adjacent sites. Sites with identical trees indicated by the same tree number (T1, T2 etc.) are grouped in grey boxes, which indicate the tree ‘span’. Note that the inferred ARG is different with unphased data because the ‘duplicable’ pattern at variant site 3 is incorrectly deemed to be compatible with that at variant site 4.

With phased haplotypes, there is only one possible suitable location for each node in the tree, and the above procedure ensures construction of the minimally resolved tree compatible with all variant patterns. In other words, every unique variant is represented by one unambiguous node, and all parent nodes that have more than two children are left as polytomies. For unphased data from higher ploidies in which some patterns are ambiguous, the heuristic procedure results in the construction of one of the possible trees compatible with the set of variant patterns. In most cases, this only affects the placement of tips from the same individual (compare for example tips B1 and B2 in the two trees in Fig. 1). However, if duplicable patterns are present, the algorithm could incorrectly infer the tree shape (by incorporating the incorrect number of duplicable patterns). An example is shown in the section on sequential tree building below.

### Collecting compatible sites

In the presence of recombination, the genealogy changes along the genome, so inference of local genealogies also involves inference of the ‘breakpoints’ that separate them. Under perfect phylogeny, there must be at least one breakpoint between every pair of variants that are not compatible with the same genealogy. Whether a pair of variants is compatible can be assessed using the four-gamete test (or a modified version that extends to any ploidy, described below). However, the challenge of building local genealogies is not as simple as finding clusters of mutually compatible sites. Under typical rates of recombination and mutation, intervals with a unique genealogy tend to be too short and contain too few mutations for inference of resolved trees from such clusters (Mailund et al. 2006; Wu 2011; Mirzaei and Wu 2017). Moreover, the decision of where to draw the breakpoints between genealogies is complicated by the fact that not all variants from distinct genealogies will be incompatible. In fact, adjacent genealogies along the genome tend to share many of the same nodes (Shipilina et al. 2023; Ignatieva et al. 2025), and this provides useful information that can inform local genealogy inference (Wu 2011; Mirzaei and Wu 2017).

Therefore, instead of identifying tracts of genome with a single genealogy and the breakpoints between them, sticcs aims to infer the local genealogy *at each variant site*. To do this, sticcs builds a matrix of compatible variant patterns at each site by ‘collecting’ compatible patterns from nearby variant sites (Fig. 2). In effect, the collecting procedure for site *i* considers each site *j* to the left and right of site *i* in order of distance and adds the pattern at site *j* to the matrix for site *i*, provided it is compatible with all other patterns already ‘collected’. In practice, for computational efficiency, sticcs avoids repeating tests of compatibility by using a ‘knockout’ algorithm that passes along the chromosome once in each direction. First, moving from left to right, a collection of compatible patterns is built by adding the pattern at each site *i* encountered and discarding (knocking out) any patterns in the existing collection that are not compatible with the pattern at site *i*. All patterns retained in the collection are then recorded as the ‘left-collection’ for site *i*. Second, a ‘right-collection’ for each site is generated using the analogous procedure, moving from right to left along the chromosome. Finally, the left- and right-collections for each site are reconciled. Incompatibilities are resolved by retaining the pattern whose nearest variant is closest to the focal site.

A downside of the knockout procedure described above is that a pattern from site *j* will only be considered for collection at site *i* if there is no variant site between *j* and *i* with a pattern incompatible with that at site *j*. This contradicts the biological reality that nodes (or ‘haplotype blocks’) are often ‘disjunct’, being shared between non-adjacent genealogies (Shipilina et al. 2023). To attempt to account for this, sticcs by default applies a ‘second chances’ rule in which a pattern is added back into the collection once the pattern that knocked it out has itself been knocked out.

### Extending the four gamete test to higher ploidy and mixed ploidies

Under the infinite sites model, two sites are not compatible with the same tree if all four haplotypes *00, 01, 10*, and *11* are present in the dataset. Here, each pair of numbers indicates whether allele 0 or 1 is present at two sites in a given haplotype. This is the four gamete test. For polarised data, we know that the ancestral haplotype *00* exists (or did in the past), so we need only observe the three derived haplotypes *01, 10*, and *11* to determine that the sites are incompatible. For unphased diploid or polyploid genotypes, we have less information, but we can still test for compatibility between sites by inferring which haplotypes are present. For diploids, the four possible haplotypes for two sites combine to produce nine possible diploid genotypes: *00, 01, 02, 10, 11, 12, 20, 21, 22*, where the two numbers indicate the number of derived alleles observed at the first and second site. Most of these provide unambigous information about which haplotypes are present. For example, diploid genotype *01* indicates the presence of haplotypes *00* and *01*, whereas diploid genotype *21* indicates the presence of haplotypes *11* and *10*. As a general rule that applies to any ploidy level, we can say that all three derived haplotypes must be present if: (i) There is at least one individual at which the number of derived alleles at the first site exceeds that at the second (implies that *10* exists), (ii) there is at least one individual at which the number of derived alleles at the second site exceeds that at the first (implies that *01* exists) and (iii) there is at least one individual at which the sum of derived alleles across the two sites exceeds the ploidy (implies that *11* exists).

This general test is exact for haploids, but for higher ploidies it makes a maximum parsimony assumption. For example, in a dataset of two individuals with genotypes *12* and *11*, we can say with certainty that the first individual carries haplotypes *01* and *11*, but we cannot say whether the second individual carries *00* and *11* or *01* and *10*. Since we cannot say conclusively that that all three derived haplotypes are present, these sites will not be deemed incompatible. The implication of this uncertainty is that it is possible for all pairs of sites in a matrix to be deemed mutually compatible, but for the complete set of patterns to not be compatible with a single genealogy (Fig. S1). In sticcs, such conflicts are resolved by incorporating nodes into the tree in order of their distance from the focal site and discarding nodes that are found to be incompatible with the tree (i.e. a suitable set of child nodes cannot be found).

### Sequential tree building

Once the collections of compatible site patterns have been generated, sticcs proceeds to tree building as described above and shown in Fig.s 1. Once the tree for the first variant site has been constructed, sticcs proceeds to the next site, moving left-to-right along the chromosome (Step 2 in Fig. 2). Because adjacent trees usually share multiple nodes, the starting point for building each subsequent site tree is not an empty tree but the tree built for the previous site (minus internal nodes that are not shared and those added multiple times in the multi-pass procedure). In addition to reducing computational time, this allows nodes to be shared between adjacent trees, allowing the inferred ARG to be represented using tskit’s ‘succinct tree sequence’ data representation (Kelleher et al. 2019; Wong et al. 2024), as well as the identification of haplotype blocks that extend across multiple trees (Shipilina et al. 2023; Ignatieva et al. 2025). Breakpoints between local trees are not directly inferred, but rather emerge during the sequential tree building whenever at least one node differs between two adjacent trees, indicating a difference in topology. By default, breakpoints are then defined as the midpoints between the last variant site assigned to the previous tree and the first variant site assigned to the next tree (Step 2 in Fig. 2).

Comparing panels A and B in Fig. 2 shows how ambiguity in unphased data can lead to errors in tree reconstruction. In haploid data (Panel A) the variants at sites 3 and 4 are incompatible, and this leads to a breakpoint between the first and second trees. In the first tree, variant patterns 1 and 3 correspond to two distinct nodes, each with three descendent leaves. However, in unphased diploid data (panel B), we can see that these are duplicate nodes with the same site pattern. Because this pattern is not deemed incompatible with that at variant site 4, and because the tree building heuristic first adds a single node for each pattern before attempting to add a second one, in this case the incorrect tree is inferred for part of the sequence. This example is included to highlight this limitation of the algorithm, but in reality duplicate nodes are rare, so this issue does not have a significant real world impact (see Results and Discussion).

### Application of sticcs for Topology weighting using the ‘stacking’ approach

As shown in the Results section, the current implementation of sticcs is most accurate for very small sample sizes, and accuracy drops rapidly with increasing sample size. This limits its usefulness for many applications. Nevertheless, the sample size limitation can be overcome during analyses that allow iteration over multiple ARGs that represent subsets of the data.

In its conventional use, topology weighting involves computation of local genealogies for a complete dataset, followed by computation of topology weights for each local genealogy (Fig. 3A). Topology weights represent the relative abundance of subtree topologies for each local tree, where each subtree has one tip representing each pre-defined group (Martin and Van Belleghem 2017). The number of possible subtree topologies depends only on the number of groups specified. Given *g* groups, there are (2 *g*−3)*!*/2^(*g*−2)^(*g*−2)*!* bifurcating rooted topologies (3 if *g*=3, 15 if *g*=4 etc.). In principle, their relative weights are computed by considering all possible subtrees in which one tip is taken from each group, and counting the number of times each topology is observed. The number of subtrees can be very large: 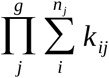, where *n*_*j*_ is the number of individuals in the *j*th group and *k*_*ij*_ is the ploidy of the *i*th individual in the *j*th group. However, in practice, the counting procedure can be sped up dramatically by accounting for redundancy in the tree (e.g. by collapsing monophyletic clades where all tips are from the same group) (Martin and Van Belleghem 2017).

**Fig. 3.**
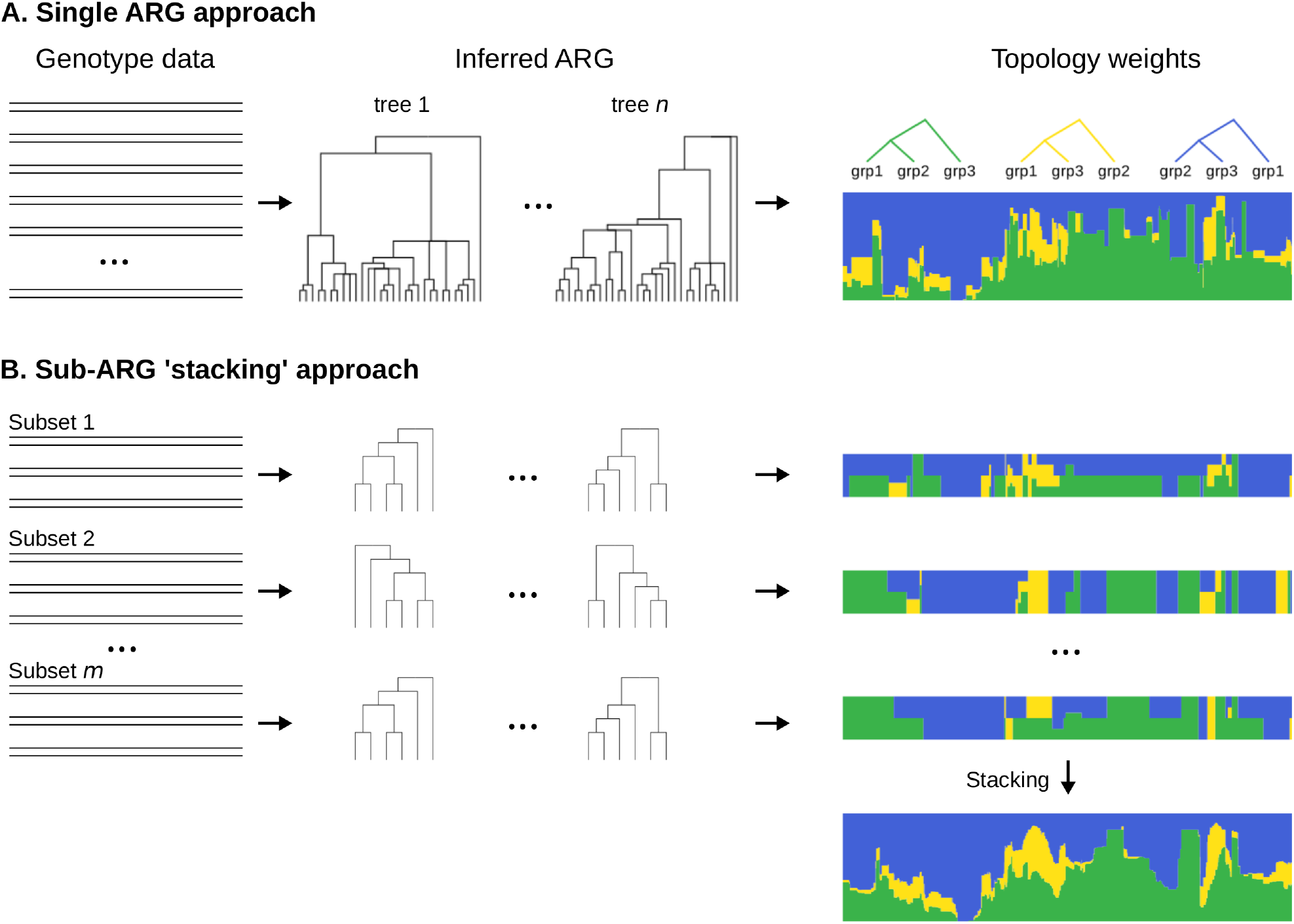
Two approaches for topology weighting starting from genotype data. **A**. In the single ARG approach, an ARG for the complete dataset is inferred and topology weights are computed for each local tree. **B**. In the ‘stacking’ approach, small sub-ARGs are inferred using subsets comprising one individual from each predefined group. Topology counts are computed for each subset and these are then stacked by summing the counts of each topology over each unique overlapping interval, resulting in the final set of topology weights. The stacking procedure allows ARG inference over small numbers of individuals, which is preferable when using sticcs.

To account for the fact that sticcs is limited to small sample sizes, we can use an alternative approach to compute topology weights: dividing the dataset into subsets of individuals, inferring a sub-ARG and counting subtree topologies for each subset, and then ‘stacking’ the results to produce the final counts (Fig. 3B). A practical minimal subset comprises a single individual from each of the *g* groups, resulting in 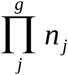 subsets. For haploid data, each subset of individuals accounts for a single subtree of each local genealogy, whereas for diploid data, each subset accounts for *2*^*g*^ subtrees. Since the tree intervals in each inferred sub-ARG will be different (because different variant patterns will be encountered with each subset of individuals), the stacking procedure involves first dividing the chromosome into the minimal set of intervals such that every breakpoint in every sub-ARG is represented and then summing the topology counts across all sub-ARGs for each interval (Fig. 3B). This stacking approach using sticcs (referred to below as sticcstack) is implemented in twisst2, along with the conventional single ARG approach (which requires a pre-inferred ARG as input).

A downside of the stacking approach is that it loses most of the computational savings achieved by simplifying redundant parts of the tree such as monophyletic clades. It will therefore be necessary in many instances to compute approximate topology weights using a limited number of subsets of individuals. Fortunately, the inferred weightings tend to converge quickly on the true values after a few hundred subtrees have been considered (Martin and Van Belleghem 2017).

An advantage of the stacking approach is that it is more robust to missing data and multi-allelic sites. Although the current implementation of sticcs does not directly handle variants with missing genotypes in some individuals, sticcstack utilises such variants for all sub-ARGs in which the subsampled individuals have non-missing genotypes. Similarly, although sticcs does not use sites with more than one derived allele, such sites are used in sticcstack for each subset in which only one of derived alleles is present (along with the ancestral allele).

### Simulation Tests

To explore the accuracy of sticcs for ARG inference and topology weighting, I simulated data using msprime (Kelleher et al. 2016; Baumdicker et al. 2022). The ‘true’ ARG was simulated using the sim_ancestry module and stored as a tree sequence object (Kelleher et al. 2016; Kelleher et al. 2019)). Variants were simulated using the sim_mutations module, and haplotypes were exported with or without further modifications. Specific simulation parameters and any downstream modifications used described along with each result. The key variations tested were finite or infinite sites mutations (the latter was simulated by identifying all sites at which recurrent mutation had occurred in the simulated tree sequence and removing all but the first mutation); different ratios of recombination rate to mutation rate (rho/theta); gene conversion; population structure; genotyping error (introducing following an empirically inferred error model (Albers and McVean 2020)) and polarisation error (added by performing inference of the ancestral state using an outgroup that split from the focal population 2N generations ago).

I first quantified the accuracy of sticcs for inferring local tree topologies and the breakpoints between them, and compared it with other popular tools: Relate (Speidel et al. 2019), tsinfer (Kelleher et al. 2019), Argweaver (Rasmussen et al. 2014) and Singer (Deng et al. 2024).To quantify accuracy using a measure that can be compared across datasets of different sizes, I computed the a normalised rooted quartet distance between the simulated and inferred local trees. This was computed as the number of quartets comprising three tips and the root node that have the same topology between the simulated and inferred tree, divided by the total number of quartets. Because the number of quartets increases very rapidly with increasing sample size, for all sample sizes over 8, I used an approximate quartet distance based on a random sample of 100 quartets. For comparing entire ARGs, this distance is computed for each unique tract of overlap between local tree intervals and then a weighted mean is taken, accounting for the span of each overlapping interval (Fig. S2). This therefore provides a compound measure of accuracy of local tree topologies and the breakpoints between them. For both Argweaver and Singer, which perform MCMC sampling of ARGs, I used a burn-in of 500 MCMC samples, and then performed the accuracy analysis on 10 sampled ARGs, with spacing of 10 for Argweaver and 20 for Singer.

Next, I explored the accuracy of each method for topology weighting specifically. To emulate a multi-species dataset, a more complex demographic model with populations that undergo divergence and admixture was used (Fig. S2). Topology weights were computed using twisst2. For Relate, tsinfer, Argweaver and Singer, the single ARG approach (Fig. 3A) was used. For sticcs, the stacking approach (Fig. 3B) called sticcstack was used, sampling 512 sub-ARGs. Accuracy was assessed by calculating the double-scaled euclidean distance between topology weights from overlapping local trees in the the simulated and inferred ARGs, weighted by the length of the overlapping tract (Martin and Van Belleghem 2017). To simulate unphased diploid and tetraploid data, haplotypes were arbitrarily combined in twos or fours and the phase information was disregarded. Based on the finding that ARG inference accuracy is primarily dependent on the ratio of recombination rate to mutation rate, three different values of rho/theta were considered: 0.1, 1 and10.

### Analysis of adaptive introgression in Heliconius butterflies

To assess the performance of sticcs on real data, I analysed previously-published data from *Heliconius* butterflies. It was recently shown that, along with adaptive introgression of alleles for mimetic wing colouration at the gene *optix*, alleles associated with mate preference at the nearby gene *regucalcin* have introgressed between the species *Heliconius melpomene* and *Heliconius timareta (Rossi et al. 2024)*. I analysed a published VCF file (Martin et al. 2019) and ran twisst2 sticcstack on chromosome 18, setting four ingroups: *H. melpomene melpomene* from French Guiana (n=10), *H. melpomene amaryllis* from Peru (n=10), *H. timareta thelxinoe* from Peru (n=10), and *Heliconius. cydno chioneus* from Panama (n=10). *Heliconius numata* (n=2) served as an outgroup for polarisation. 512 sub-ARG samples were taken.

## Data Availability Statement

The sticcs Python package is available at https://github.com/simonhmartin/sticcs. The twisst2 Python package is available at https://github.com/simonhmartin/twisst2. Scripts for running all tests with simulated and real data and plotting the results are available at https://github.com/simonhmartin/testing-sticcs-twisst2. All other data necessary for confirming the conclusions of the article are present within the article.

## RESULTS

### ARG inference

sticcs infers the topology and span of each local genealogy along the genome, and does not attempt to infer coalescence times. I therefore used the normalised quartet distance to compare topologies between overlapping local trees between the ‘true’ and inferred ARGs (see Methods for details). When sample size is very small (n=4), sticcs is more accurate than existing methods for ARG inference under most simulated conditions (Fig. S3). However, while the other methods maintain similar accuracy with increasing sample size, the accuracy of sticcs drops. By n=32, sticcs is less accurate than all other tested methods. One exception is under strong population structure, where all the methods perform similarly well (Fig. S3G). The drop in accuracy for sticcs in panmictic populations is seen under both finite-sites and infinite sites mutation simulations (Fig. S3A, S3B), indicating that it is not simply a result of violation of the assumptions of perfect phylogeny. Instead, the drop in accuracy is associated with an increase in unresolved nodes (Fig. S3). This indicates that, under large sample sizes, the heuristic procedure for collecting compatible site patterns from further afield begins to fail, leading to a lack of variant information to fully reconstruct each local tree. According to the metric used, Relate has the best accuracy for topology inference at larger sample sizes.

Increasing the recombination rate to mutation rate ratio (rho/theta) from 1 to 10 causes a strong reduction in accuracy for all methods (Fig. S3C). It also worsens the relationship between sample size and accuracy for sticcs, and the number of polytomies increases dramatically. This shows that the limitations of the site collecting heuristic with increasing sample size are particularly pronounced when rho/theta is high. Adding realistic gene conversion has a comparatively minor impact on accuracy (Fig. S3D). Adding realistic genotyping error (Fig. S3E) and polarisation error (Fig. S3F) in the simulation causes a small decrease in accuracy of a similar magnitude for all methods. Across all simulated conditions, only Argweaver and Singer tend to infer accurate numbers of trees (changes of topology along the chromosome), but this is expected as they were provided with the recombination rate, and this is incorporated in the inference model. Relate and sticcs tend to underestimate the true number of trees by the greatest margin, despite showing high overall accuracy in ARG topology inference. This indicates that the ‘missed’ trees probably tend to be very narrow and/or very similar in topoology to those around them.

At small sample sizes (16 or below), sticcs is the fastest of the methods tested, followed by Relate, tsinfer, Singer and Argweaver, which is orders of magnitude slower (Fig. S4). However, sticcs scales poorly with sample size, with nearly quadratic time complexity. It scales linearly with sequence length, and this depends on the number of variants rather than the number of recombinations underlying the simulated data (Fig. S4).

### Topology weighting

Despite the fact that the sticcs algorithm is only useful for small sample sizes, it can be used for topology weighting with any sized dataset using the stacking approach (‘sticcstack’) described in the Methods section and Fig. 3B, and implemented in twisst2. Topology weighting was performed on datasets simulating species diverging and admixing, with either three or four ingroups (Fig. S5). Of the methods tested, sticcstack tends to have the highest accuracy, especially at fine resolutions (Fig. 4A). In general, all methods perform better when the rho/theta ratio is low, reflecting the greater information content, but the different methods do not respond equally. Accuracy of topology weights also improves when they are averaged over larger genomic regions, again with subtle differences in the shape of the relationship for different methods (Fig. 4A). Remarkably, sticcstack shows almost no drop in accuracy when provided with unphased diploid data, except when rho/theta is very low (Fig. 4A). sticcstack run with unphased tetraploid data shows lower accuracy, but it is comparable with that of the other methods applied to perfectly phased data. Despite the measurable differences in the quantified accuracy, visual comparison shows that all of the tested methods capture topology weights accurately and with similarly high resolution (Fig. 4B, Fig. S6, S7).

**Fig. 4.**
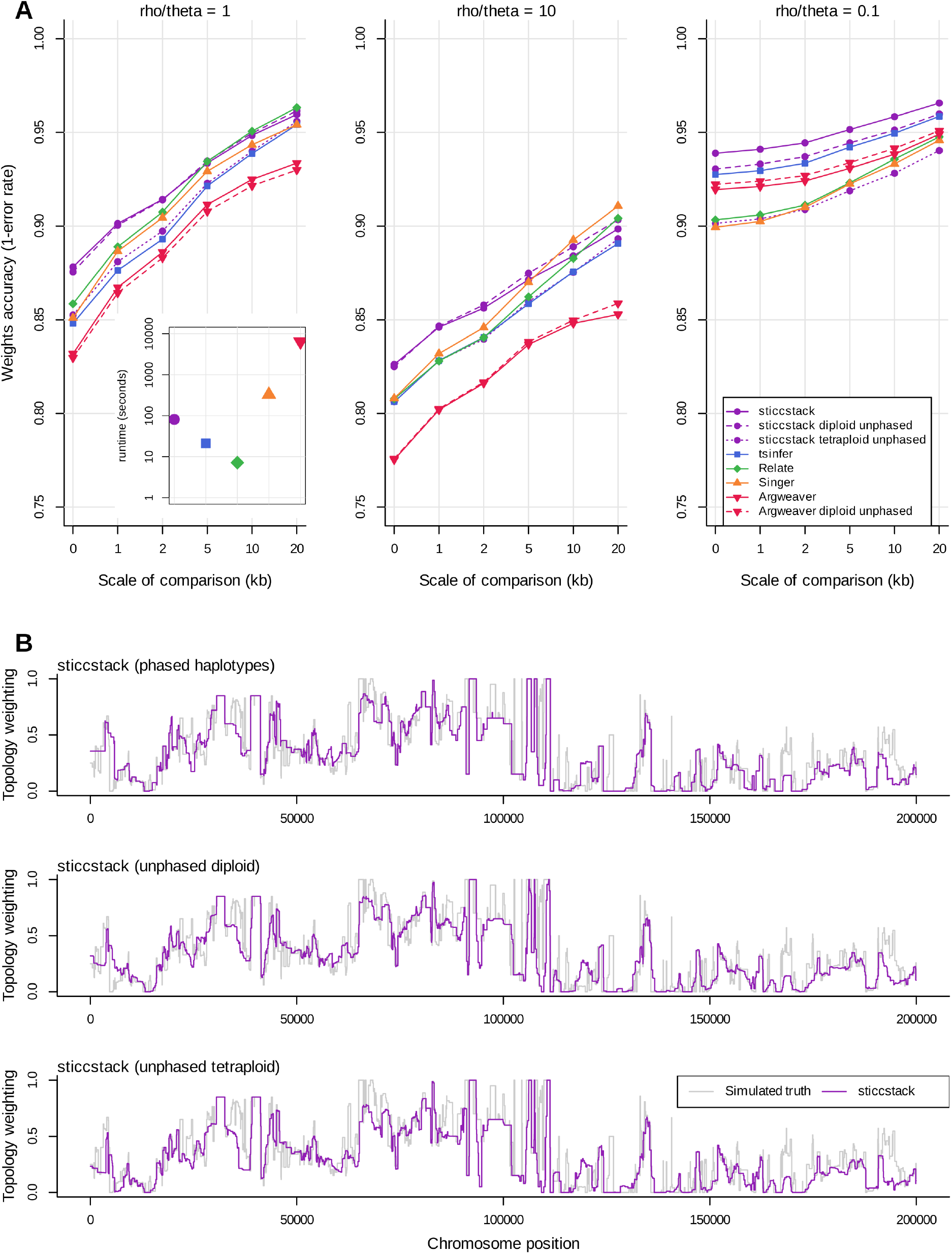
Comparison of topology weighting accuracy. **A**. Accuracy of each method in the four-species admixture simulation. X-axis indicates the level of smoothing by averaging over a genomic window, with 0 indicating no smoothing. Each point represents the average accuracy over 40 simulations of a 100 kb region. Panels represent different rho/theta values. The insert indicates the average runtime for each analysis of a 100kb region. **B**. sticcstack inferred weights one of the topologies in the three species admixture simulation compared to the simulated truth (in grey).

Because sticcstack uses a set sample size for each sub-ARG, its runtime depends linearly on the number of sub-ARGs to be sampled (and the length of the sequence). For the tests described here I sampled 512 sub-ARGs. The runtime for the full topology weighting analysis was slower than for Relate and tsinfer, but four times faster than Singer and 80 times faster than Argweaver (Fig. 4A insert).

### Analysis of adaptive introgression in Heliconius butterflies

Of the 15 possible topologies describing the relationships between the analysed *Heliconius* species, estimated weightings confirmed the most abundant across chromosome 18 is the accepted species tree (Fig. 5A, 5B). A topology consistent with introgression between *H. melpomene amaryllis* and *H. timareta thelxinoe* shows several peaks near the chromosome ends. The largest two peaks correspond to the regulatory region near *optix* known to control variation in red pigmentation (Wallbank et al. 2016) and *regucalcin*, recently shown to contribute to mate preference (Rossi et al. 2020; Rossi et al. 2024). These patterns are clearer than those observed using window-based trees (Rossi et al. 2024), and further strengthen the evidence for co-introgression of wing colouration and visual mate preference alleles between *Heliconius* species.

**Figure 5.**
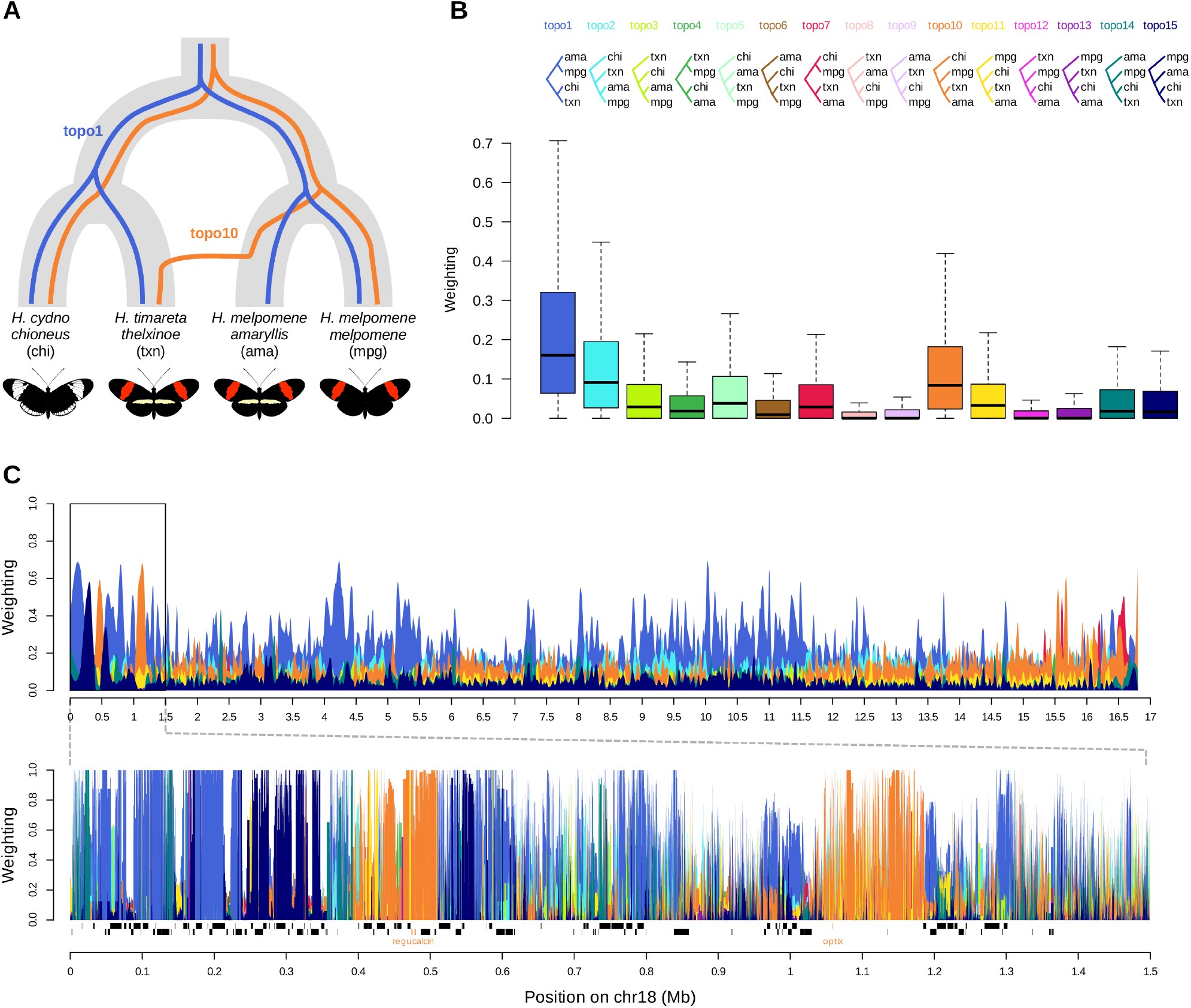
Analysis of adaptive introgression signals in Heliconius butterflies. **A**. The accepted species tree for the four populations of interest. Superimposed are two subtree topologies of interest: topo1 (the species topology) and topo10 (consistent with introgression from H. melpomene amaryllis into H. timareta thelxinoe). **B**. Boxplots of weightings for all 15 possible topologies across chromosome 18. **C**. Topology weightings from sticcstack plotted across chromosome 18 with smoothing to show overall trends. **D**. Raw topology weightings from sticcstack plotted across the 1.5 Mb region of interest. Genes are shown along the bottom of the plot, with the two genes of interest, regucalcin and optix in orange.

## DISCUSSION

The primary motivation for developing sticcs was to explore the possibility of genealogical inference without attempting to phase haplotypes, especially for the purpose of topology weighting studies. Phase inference from short read sequencing data usually relies on patterns of linkage-disequilibrium, making assumptions about (lack of) population structure, and ideally using reference panels of phased haplotypes (Delaneau et al. 2019; Browning et al. 2021). These approaches are not well suited to small multi-species datasets from non-model taxa, especially polyploids. Importantly, many tree-based analyses, including topology weighting, only require that tips are labelled by the individual or group they represent, making phased haplotypes theoretically unnecessary. I have shown that, at least for small sample sizes, an ancestral recombination graph (ARG) can be reliably inferred from unphased data using the principles of perfect phylogeny, without a phasing step. Although sticcs does considers the haplotypes from each unphased individual as interchangeable, by ensuring consistency of shared nodes across adjacent trees, it does actually perform a sort of short-range phase inference. However, this is not intended as a substitute for long-range haplotype reconstruction. One currently available tool for ARG inference, ARGweaver, can analyse unphased diploid data (Rasmussen et al. 2014; Hubisz et al. 2020), but this is much slower and relies on assumptions about the underlying demography. Unlike ARGweaver and several other tools, sticcs reconstructs only the topology of local genealogies, and does not attempt to infer coalescence times. In principle these could be inferred for an ARG output by sticcs using for instance tsdate (Wohns et al. 2022).

The major shortcoming of the sticcs algorithm as described here is that it is only comparable in accuracy with existing ARG inference methods when sample size is very small (around 16 haplotypes or less). Contrary to expectations, this is not primarily a result of violating the perfect phylogeny assumptions under recurrent mutation and genotyping errrors. A similar decline in accuracy is seen under finite and infinite-sites simulations, and with and without genotyping errors. Instead, the heuristic algorithm for collecting information about tree nodes from further afield on the chromosome appears to be inherently limited with increasing sample size. Haplotype blocks (i.e. groups of lineages that share a common ancestor for a given stretch of the genome), can be disjunct, appearing in non-adjacent local genealogies (Shipilina et al. 2023). By discarding site patterns that are separated by too many incompatible sites, the sticcs algorithm fails to fully capture information shared by disjunct haplotype blocks when the sample size is large. This issue may be addressed in the future with modifications to the algorithm, but probably at some computational cost.

Fortunately, for purposes like topology weighting, where the problem can be broken down across many small sub-ARGs, the sample size limitation of sticcs can be circumvented. Topology weights estimated using the stacking approach implemented in twisst2 (sticcstack) are more accurate than those computed from full ARGs inferred using Relate (Speidel et al. 2019), tsinfer (Kelleher et al. 2019), Singer (Nielsen et al. 2025)or Argweaver (Rasmussen et al. 2014). sticcstack remains accurate when applied to unphased diploid data, and has acceptable accuracy when applied to unphased tetraploids. There is, however a computational cost to the stacking approach. When computing topology weights from a single genealogy, the large combinatorial problem can be simplified by effectively collapsing redundant parts of the tree (Martin and Van Belleghem 2017). By splitting the weighting computation over many inferred sub-ARGs, these computational savings are lost. For this reason, it will often be necessary to compute approximate topology weights based on a non-exhaustive set of sub-ARGS inferred from randomly sampled individuals. Fortunately, the estimated topology weights converge quickly towards the truth after a few hundred subtrees have been sampled (Martin and Van Belleghem 2017).

The intrinsic ambiguity in unphased data means that there is less information available for tree building. In addition to the fact that tips from the same individual are considered interchangeable (which does not impact topology weighting), an incorrect tree can be reconstructed if the algorithm fails to incorporate the correct number of ‘duplicate’ nodes with the same site pattern (see Methods and Fig. 2B). However, the high level of accuracy observed in tests of sticcstack with unphased data shows that this issue does not pose a major problem in realistic data. Indeed, in simulated genealogies for four diploid individuals, only 3.5% of nodes are duplicate nodes, and this drops to 1.5% under the multi-spcies admixture model analysed here. Reduced accuracy in analysing unphased tetraploids is therefore probably more a result of the unavoidably increased sample size of each sub-ARG being inferred (as there are four haplotypes per sampled individual), and the consequent issues described above, than of the unphased data *per se*. Future improvements to the collecting algorithm may therefore further improve performance on unphased data.

Most studies that have applied topology weighting to date have used crude window-based tree inference. This approach is undesirable not only because it fails to use information from outside the focal interval for tree reconstruction, but because it does not attempt to identify breakpoints between local trees, and thereby will tend to average over genomic regions with distinct genealogies. Localising the specific genomic region with a distinct topology weighting is particularly desirable when attempting to identify loci associated with trait variation or adaptive introgression (Belleghem et al. 2017; Marburger et al. 2019; Morris et al. 2019; Stankowski et al. 2023; Rossi et al. 2024), which can be extremely narrow (Marburger et al. 2019). The sticcstack method was recently used to identify narrow tracts of ‘gene flux’ between inversion haplotypes, associated with a change in wing colouration in the butterfly *Danaus chrysippus* (De-Kayne et al. 2025).

Any method for fine-scale ARG inference will be inherently limited by the ratio of recombination rate to mutation rate (rho/theta). While this ratio is on average around 1 in humans, it can be >10 in recombination hotspots, or even genome wide in taxa such as butterflies (Keightley et al. 2015; Davey et al. 2016), meaning that many narrow genomic regions with distinct genealogies will tend to be unidentifiable due to a lack of mutational information. It is important to note that this problem cannot be solved by the common practice of window-based tree inference over large blocks of hundreds or thousands of SNPs in an attempt to reconstruct an ‘average’ tree (Martin et al. 2013). In fact, forcing tree-like evolution on a large recombinant region has the opposite to the desired effect, causing a bias towards the most abundant topology in the genome and drowning out the signal of less abundant local topologies (Martin and Van Belleghem 2017). An interesting recent alternative approach to topology weighting uses deep learning to infer topology weights directly from unphased sequencing data without the need to infer local trees (Y. Zhang et al. 2023). This approach, called ERICA, is designed to detect introgression, but in doing so it infers the *average* topology weights across genomic windows, assuming that a considerable amount of recombination has occurred within each window. It therefore actually *increases* in accuracy with increasing recombination rate, but it only captures average signals, and does not provide fine-scale resolution of how topologies change along the genome.

A valuable characteristic of MCMC sampling methods for ARG inference like Argweaver and Singer is that they provide information on uncertainty, and therefore on the information content of the underlying data. This distinguishes them from methods that rely on heuristics, like tsinfer, Relate and sticcs, but it comes at the cost of greater reliance on underlying model assumptions (though this did not appear to be a major impairment in the ‘multi-species’ simulation scenarios tested here). One advantage of the particularly simple algorithm used by sticcs is that its output is ‘close to the data’. In this way, it is similar to the intuitive ‘ABBA-BABA’ methods that use mutation patterns as proxies for the topologies of underlying genealogies (Durand et al. 2011). It may therefore be useful to think of the ARGs inferred by sticcs and topology weights computed using sticcstack as lying somewhere between an informative summary of the dataset and an actual inference of the evolutionary process that produced it.

## ACKNOWLEDGEMENTS

This work was funded by a fellowship from the Royal Society (grant codes URF\R1\180682 and URF\R\231034). Thank you to Konrad Lohse, Alexander Mackintosh and Jerome Kelleher for helpful input.

