## Supplementary Figures for "A model-free method for genealogical inference without phasing and its application for topology weighting"

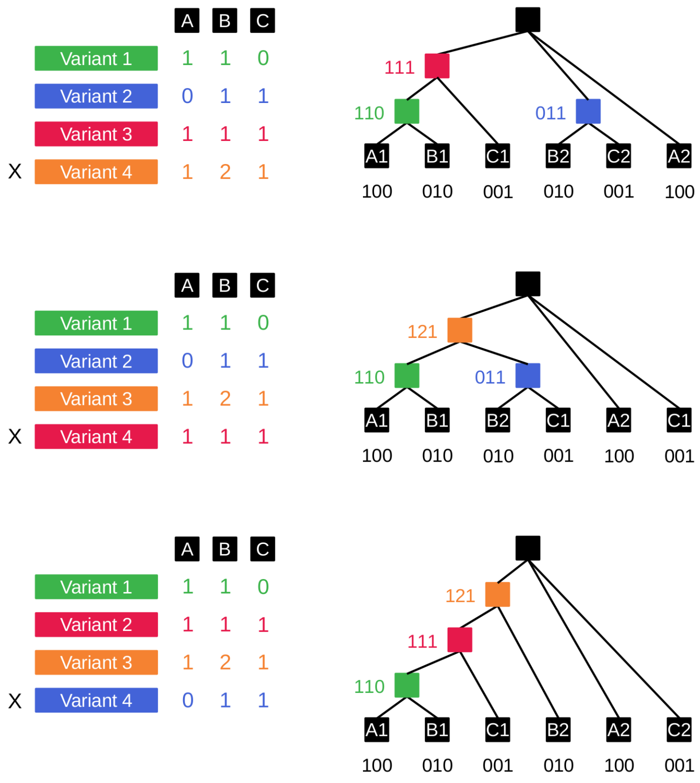


***Fig. S1. Examples of conflicts when applying perfect phylogeny tree building to diploid genotypes.*** *Four variant patterns for three diploid individuals are shown, in three different orderings. All pairs of variant patterns are mutually compatible according to the generalised four gamete test described in the main text. Nevertheless, the four variant patterns are not compatible with a single tree. Applying the ordered tree building procedure shown in Fig. 1 results in a different final tree depending on the order in which variant patterns are added. In in each case one pattern is discarded (indicated by 'X') because no suitable child nodes can be found.*

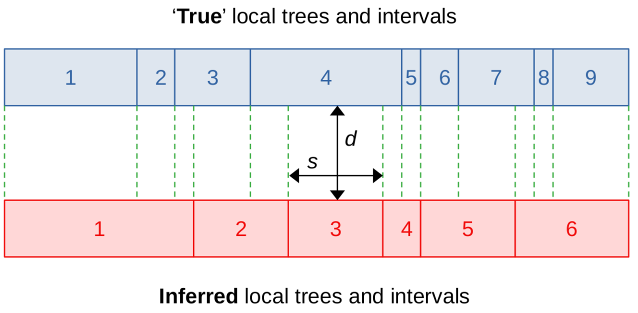


$$D = \frac{\sum^{n} d_i s_i}{\sum^{n} s_i}$$

***Fig. S2. Diagram showing how ARG inference accuracy is quantified.*** *The ARG can be represented as a series of local trees separated by breakpoints. Here each numbered box indicates a different local tree and its span on the chromosome. Inferred ARGs can differ from the true underlying ARG not only in their tree topologies, but also in the number of trees and the locations of breakpoints. To compare topologies in an inferred ARG (red) to the truth (blue), we therefore first need to decide which trees to compare with which. This is done by dividing the chromosome at every breakpoint in both ARGs (dashed lines). This creates a new set of n intervals. Each of the n intervals overlaps exactly one tree in the true ARG and one tree in the inferred ARG. We therefore compute n distances (d), one for each of the n intervals. To get the overall distance (D) we take a weighted mean of the n values of d, weighted by the relative span (s) of each interval.*

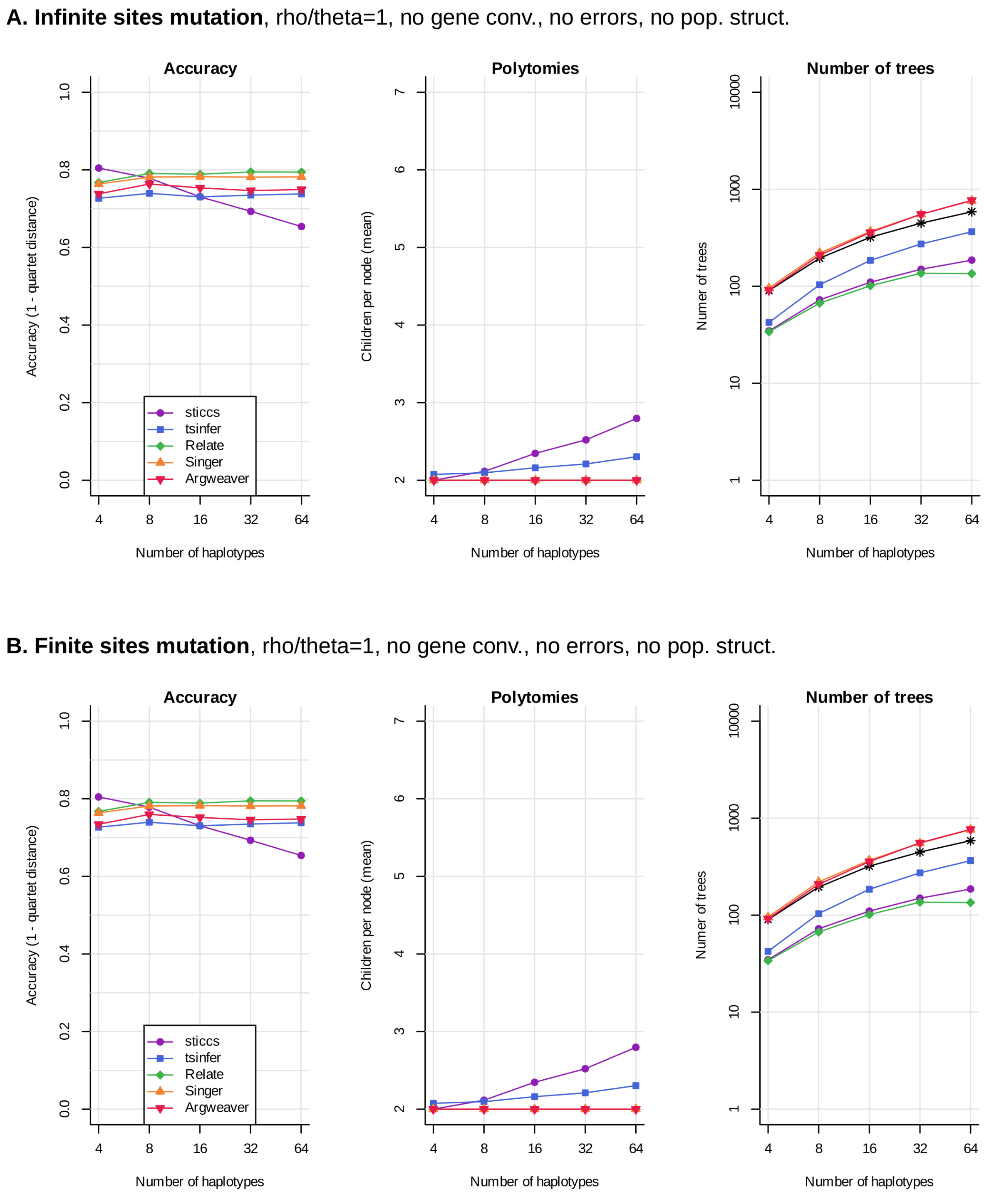


***Fig. S3. Assessment of ARG inference accuracy (panels A and B).*** *Left panel shows the accuracy of local tree topology and breakpoint inference using sticcs and four other popular ARG inference tools. The x-axis indicates the number of haplotypes analysed. Middle panel shows the mean number of children per node per tree, with deviations above 2.0 indicating the presence of unresolved nodes (polytomies). Note that only sticcs and tsinfer allow polytomies. Right panel shown the total number of trees in the 100kb ARG inferred using each method, with the simulated true number shown with black astrices. All plotted results are averaged over 20 simulations of a 100kb region.* ***A****: infinite sites mutational model, $N_e$=1e5, mu=1e-8, r=1e-8.* ***B****: Finite sites mutational model, $N_e$=1e5, mu=1e-8, r=1e-8.*

**C.** Finite sites mutation, **rho/theta=10**, no gene conv., no errors, no pop. struct.

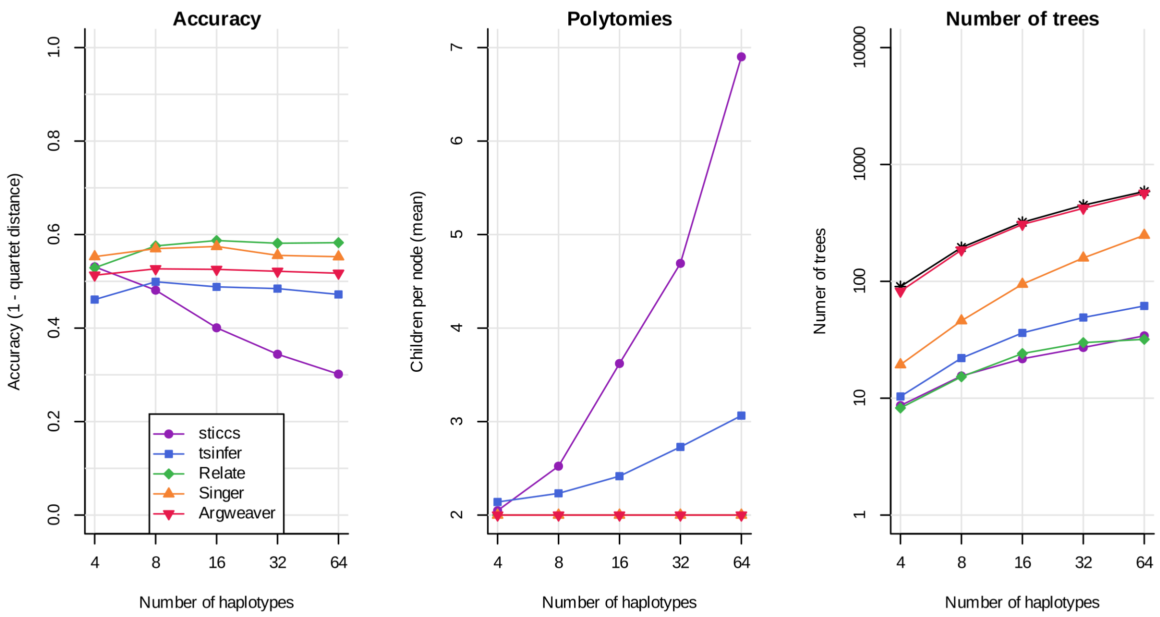


**D.** Finite sites mutation, rho/theta=1, **with gene conv.**, no errors, no pop. struct.

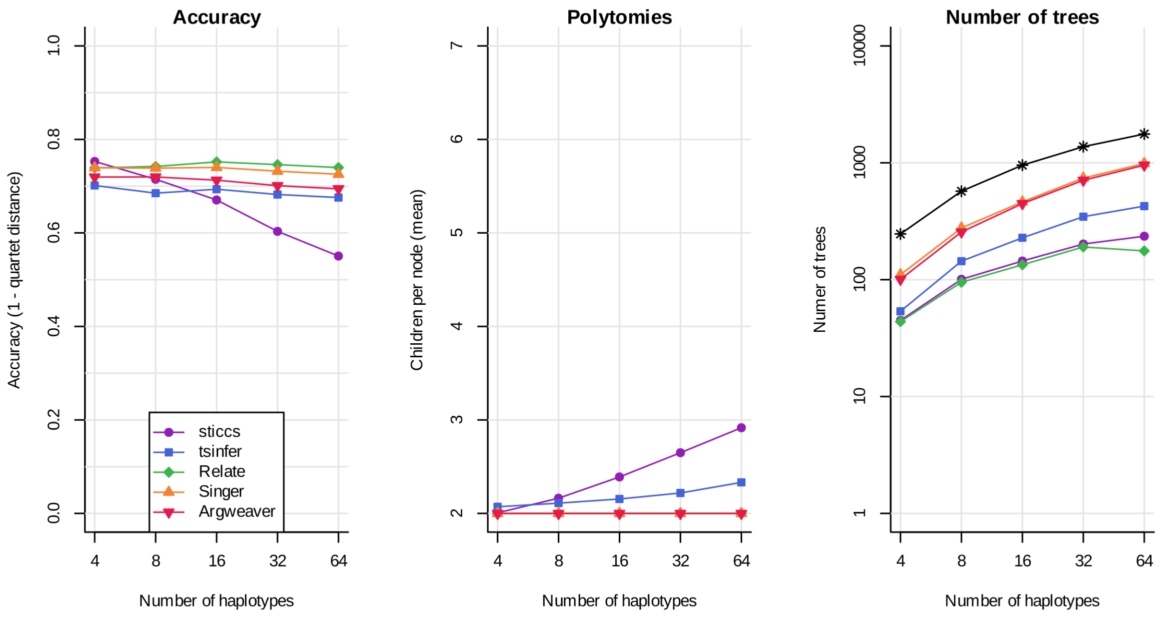


***Fig. S3. Assessment of ARG inference accuracy (panels C and D).*** *Left panel shows the accuracy of local tree topology and breakpoint inference using sticcs and four other popular ARG inference tools. The x-axis indicates the number of haplotypes analysed. Middle panel shows the mean number of children per node per tree, with deviations above 2.0 indicating the presence of unresolved nodes (polytomies). Note that only sticcs and tsinfer allow polytomies. Right panel shown the total number of trees in the 100kb ARG inferred using each method, with the simulated true number shown with black astrices. All plotted results are averaged over 20 simulations of a 100kb region.* ***C****: with high recombination rate,* $N_e$*=1e5, mu=1e-9, r=1e-8.* ***B****: with gene conversion,* $N_e$*=1e5, mu=1e-8, r=1e-8, GC rate=1e-8, GC mean tract length = 300.*

**E.** Finite sites mutation, rho/theta=1, no gene conv., **with genotyping errors**, no pop. struct.

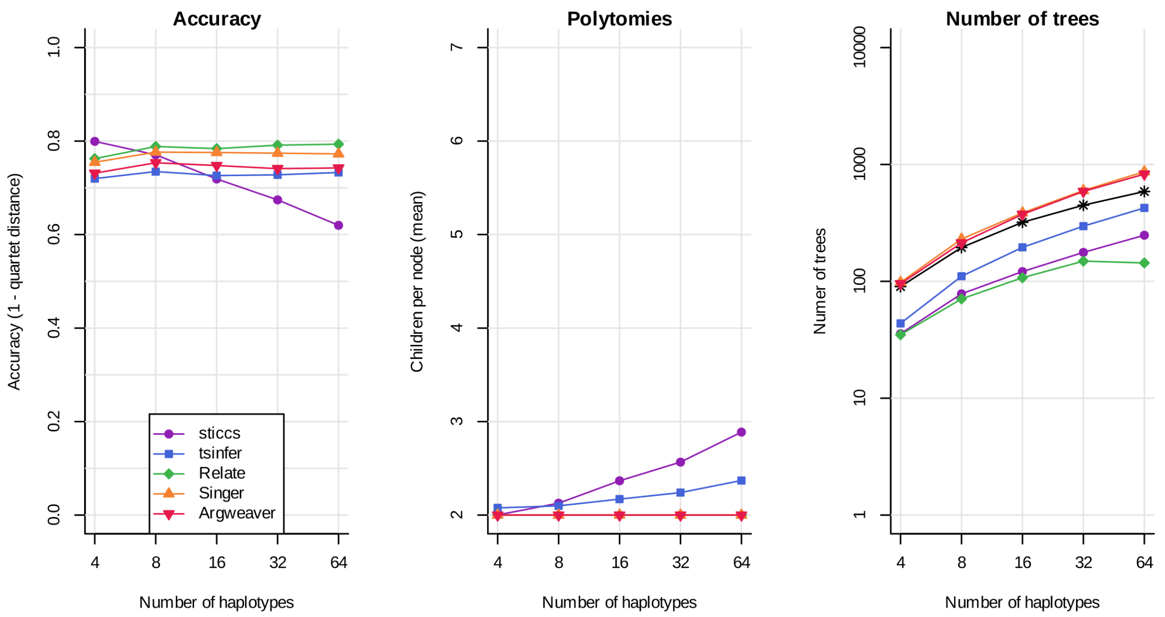


**F.** Finite sites mutation, rho/theta=1, with gene conv., **with polarization errors**, no pop. struct.

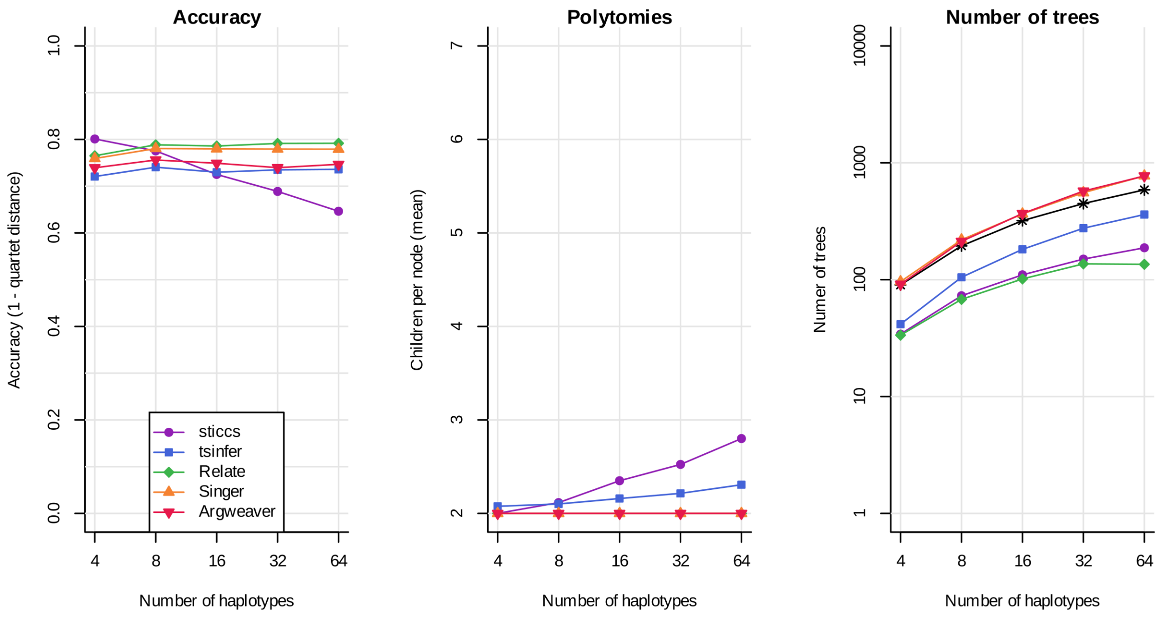


***Fig. S3. Assessment of ARG inference accuracy (panels E and F).*** *Left panel shows the accuracy of local tree topology and breakpoint inference using sticcs and four other popular ARG inference tools. The x-axis indicates the number of haplotypes analysed. Middle panel shows the mean number of children per node per tree, with deviations above 2.0 indicating the presence of unresolved nodes (polytomies). Note that only sticcs and tsinfer allow polytomies. Right panel shown the total number of trees in the 100kb ARG inferred using each method, with the simulated true number shown with black astrices. All plotted results are averaged over 20 simulations of a 100kb region.* ***E:*** *with genotyping errors, $N_e$=1e5, mu=1e-8, r=1e-8.* ***B:*** *with polarisation errors, $N_e$ =1e5, mu=1e-8, r=1e-8.*

**G.** Finite sites mutation, rho/theta=1, no gene conv., no errors, **with pop. struct. (Fst ~ 0.5)**

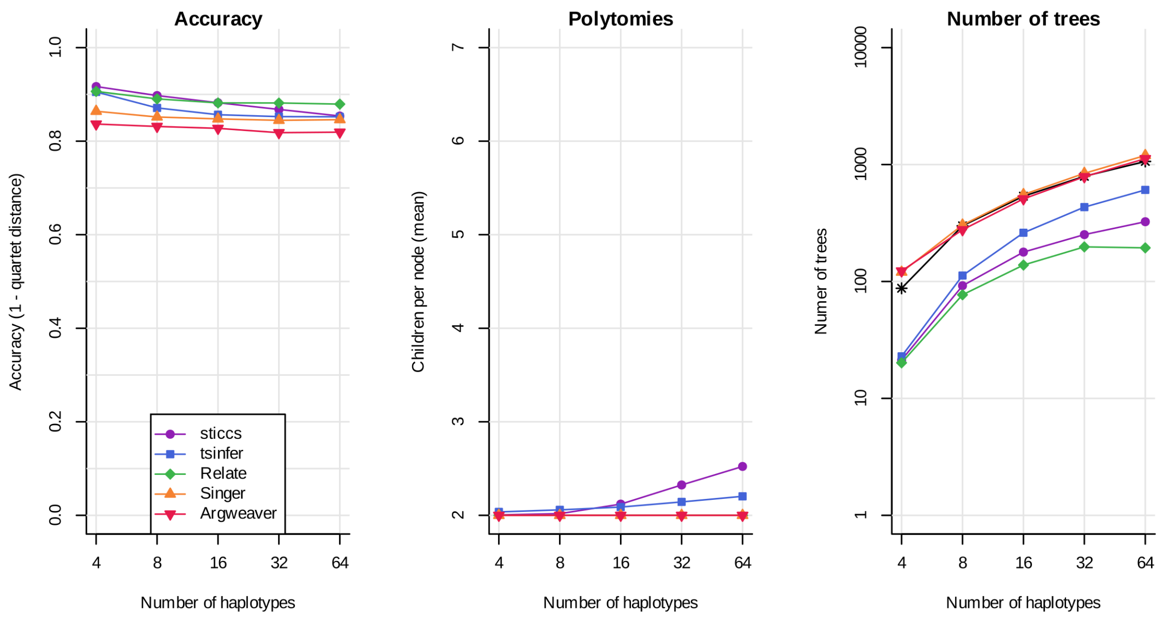


***Fig. S3. Assessment of ARG inference accuracy (panel G).*** *Left panel shows the accuracy of local tree topology and breakpoint inference using sticcs and four other popular ARG inference tools. The x-axis indicates the number of haplotypes analysed. Middle panel shows the mean number of children per node per tree, with deviations above 2.0 indicating the presence of unresolved nodes (polytomies). Note that only sticcs and tsinfer allow polytomies. Right panel shown the total number of trees in the 100kb ARG inferred using each method, with the simulated true number shown with black astrices. All plotted results are averaged over 20 simulations of a 100kb region.* ***G****: with population structure, $N_e$=1e5, mu=1e-8, r=1e-8, split time = 2N generations ago.*

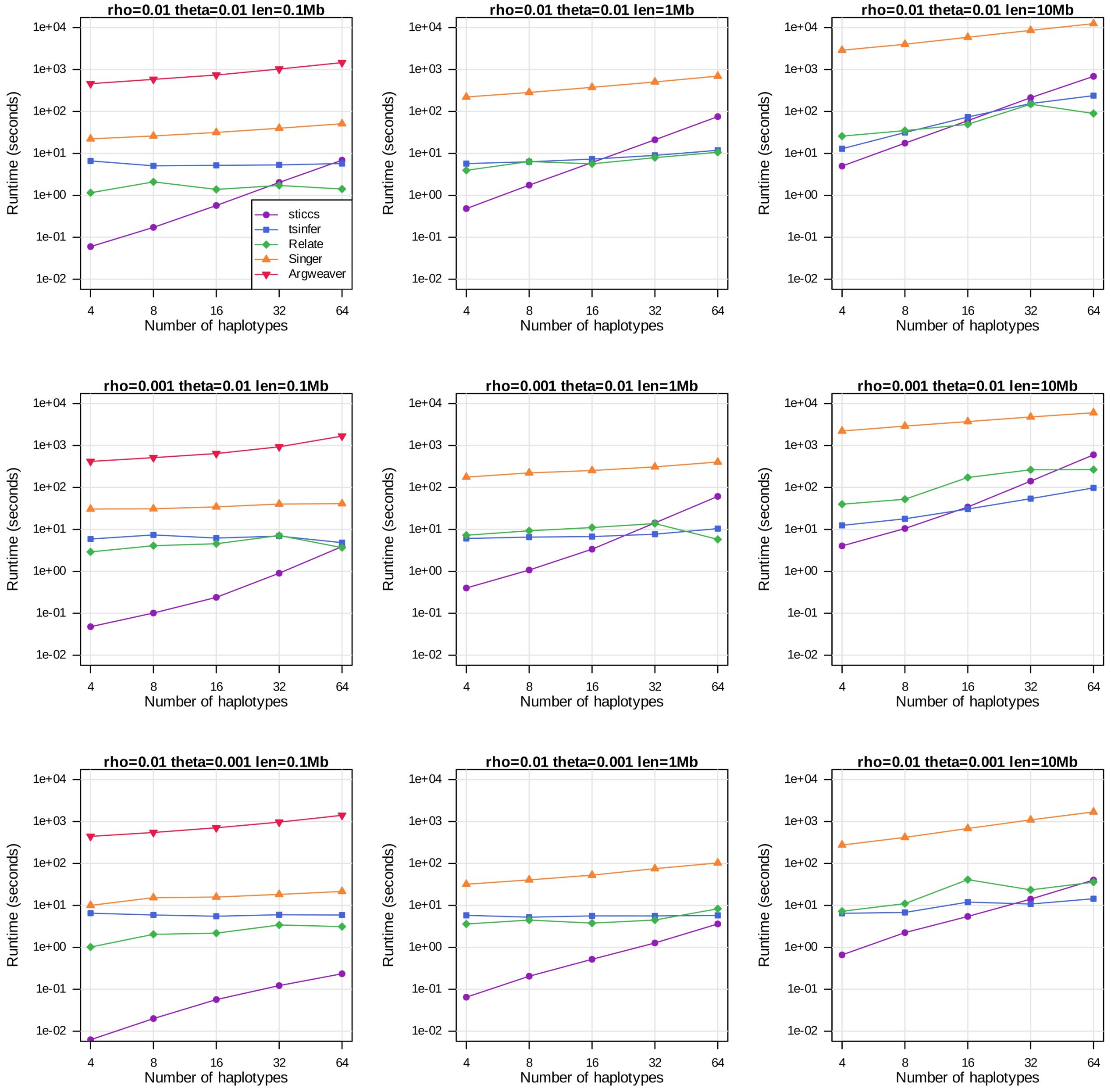


***Fig. S4. Runtime comparisons.*** *Each panel shows the runtime of each method for a given rho=$N_e r$, theta= $N_e \mu$ and sequence length. The runtime of sticcs is nearly quadratic with sample size. It is linear with sequence length (equivalent points in graphs of the same row jump by an order of magnitude as the sequence is increased by the same amount). Runtime is slowest when theta is high (first tow rows), and does not depend much on rho. Argweaver was only tested at the 0.1Mb scale due to its slow runtime.*

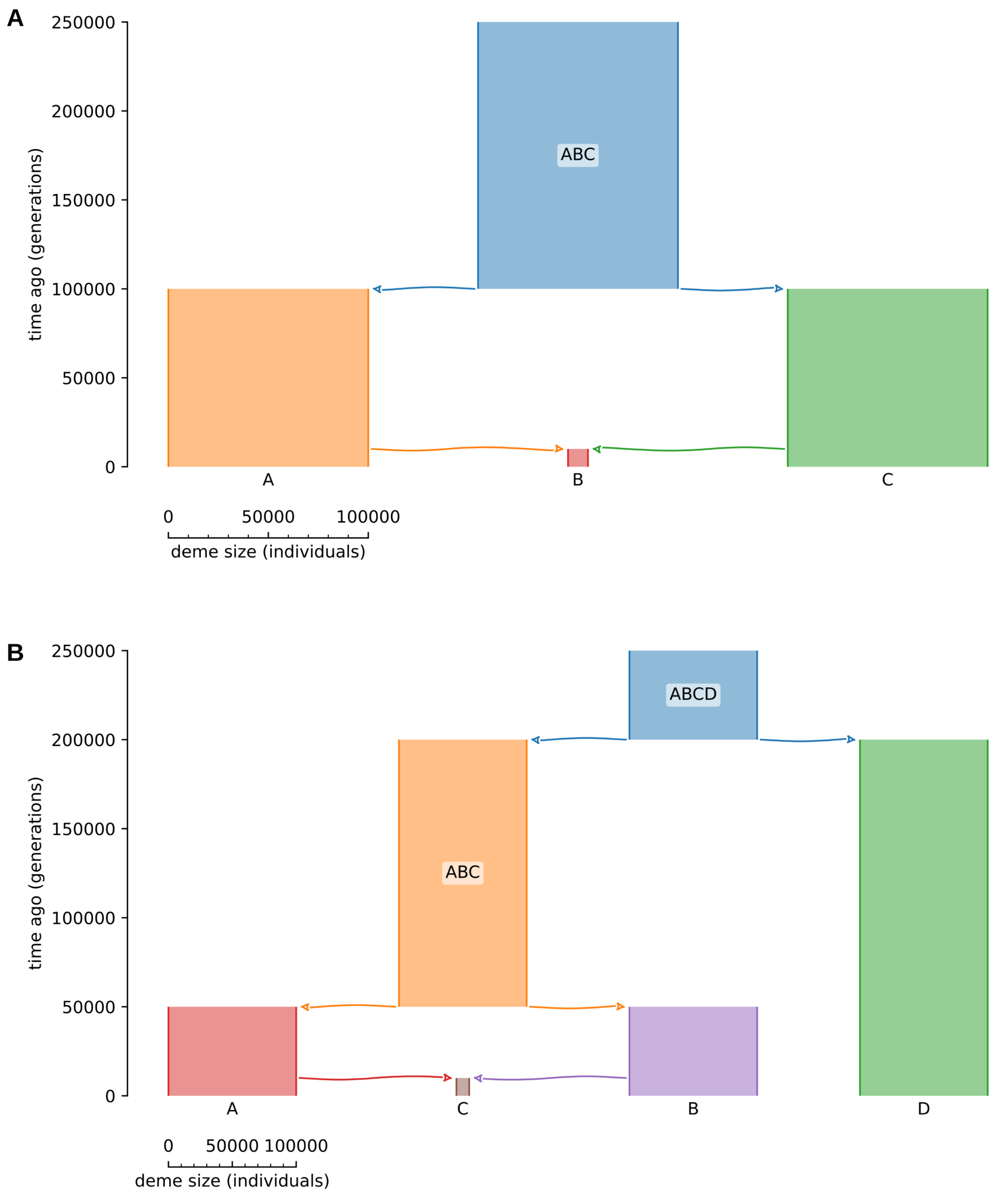


***Fig. S5. Simulation models for topology weighting tests. A**. With three ingroups. **B**. With four ingroups. Scenarios of admixture in a small recipient population were used because this causes large fluctuations in ancestry along the genome without the need to simulate selection. Equal admixture in both directions was simulated.*

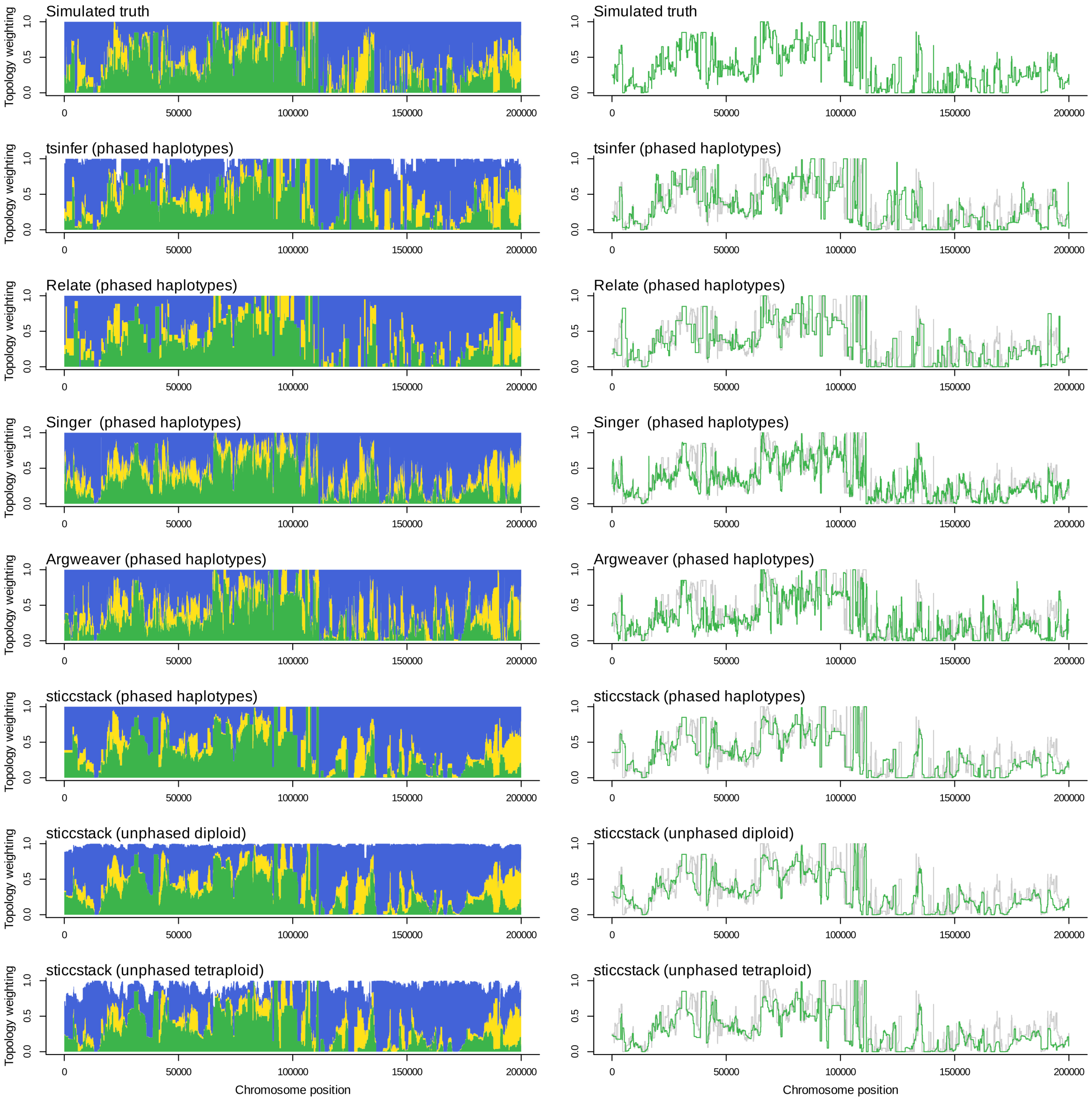


***Fig. S6. Visual comparison of topology weights inferred using different methods.*** *Simulations were performed under the three-species admixture scenario (Fig. S5A). The top row shows the simulated truth. Subsequent rows show weightings computed using inferred ARGs using different methods. Left panels show weightings for all three subtree topologies. Right panel shows weightings for one of the topologies plotted over the simulate truth in grey.*

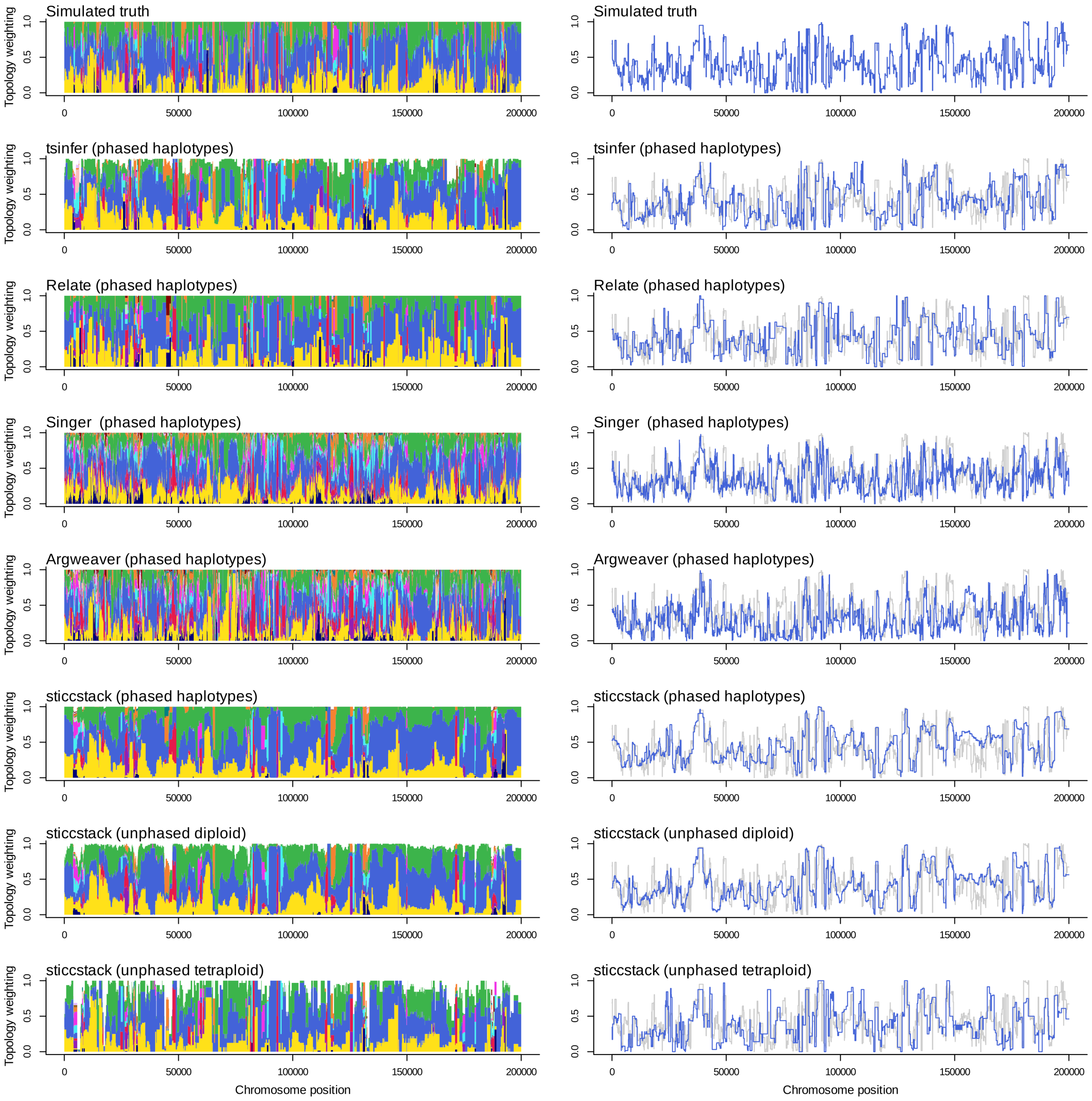


***Fig. S7. Visual comparison of topology weights inferred using different methods.*** *Simulations were performed under the four-species admixture scenario (Fig. S5B). The top row shows the simulated truth. Subsequent rows show weightings computed using inferred ARGs using different methods. Left panels show weightings for all three subtree topologies. Right panel shows weightings for one of the topologies plotted over the simulate truth in grey.*
